# Revisiting face-to-hand area remapping in the human primary somatosensory cortex after a cervical spinal cord injury

**DOI:** 10.1101/2025.03.11.642631

**Authors:** Paige Howell, Finn Rabe, Charlotte Meneghin, Simon Schading-Sassenhausen, Hunter Schone, Harshal Arun Sonar, Jamie Paik, Patrick Freund, Nicole Wenderoth, Sanne Kikkert

## Abstract

Spinal cord injury (SCI) leads to profound disruptions in sensorimotor processing. Seminal research in non-human primates suggests this sensory deprivation causes functional remapping in the primary somatosensory cortex (S1), where somatotopic representations of deprived body parts, such as the hand in cervical SCI, become responsive to touch on intact body parts, such as the face. However, evidence for such remapping in humans remains inconclusive.

We investigated face-to-hand remapping in 16 chronic cervical SCI patients and 21 able-bodied controls using two fMRI experiments. Experiment 1 employed a lip movement task, while Experiment 2 investigated the full architecture of S1 face reorganisation through vibrotactile stimulation of the forehead, lips and chin. We assessed (1) the level of face activity in the anatomical S1 hand area, (2) cortical shifts in peak face activity, (3) face-part separability in the S1 hand area and (4) correlations with clinical characteristics that may drive face-to-hand area remapping. Our results revealed no significant evidence in favour of face-to-hand area remapping in tetraplegic patients across markers of face-to-hand remapping during either lip movement or vibrotactile stimulation of face parts. Furthermore, our markers of remapping did not correlate with clinical characteristics.

These findings suggest that cortical face-to-hand remapping is not apparent in human cervical SCI patients. Beyond providing insight into the limitations of cortical reorganisation in humans, this highlights the need to reassess rehabilitation strategies based on the assumption of large-scale face-to-hand reorganisation after an SCI.

## 1. Introduction

A spinal cord injury (SCI) profoundly alters the sensorimotor system, disrupting both muscle function and sensation in the limbs and torso below the level of injury (McDonald & Sadowsky, 2002). Seminal non-human primate studies demonstrate that this loss of sensation leads to extensive remapping in brain areas containing detailed map-like body representations, such as the primary somatosensory cortex (S1) (Jain, Catania, & Kaas, 1997; Jain, Florence, & Kaas, 1998; Kaas et al., 2008). Consequently, cortical areas that are deprived of sensory inputs, such as those from the hand, become responsive to touch on intact body parts, such as the face (Reed, Liao, Qi, & Kaas, 2016). While animal models of SCI have consistently demonstrated remapping in S1 (Jain et al., 1997; Jain, Qi, Collins, & Kaas, 2008; Kambi et al., 2014; Pons et al., 1991), evidence for similar remapping following human SCI remains inconclusive (see Melo, Macedo, and Soares (2020) for a review).

Several studies have demonstrated evidence in favour of S1 remapping after human SCI, where activation from touch or movement of intact body parts is observed in or shifted towards the deprived S1 area (Bruehlmeier et al., 1998; Corbetta et al., 2002; Freund et al., 2011; Moore et al., 2000; Perani et al., 2001); though see (Henderson, Gustin, Macey, Wrigley, & Siddall, 2011; Turner, Lee, Schandler, & Cohen, 2003; Wrigley et al., 2009). However, most of these studies were done in thoracic or lumbar SCI patients, in whom sensorimotor inputs to the cortical hand area, adjacent to the face representation, remain intact (Henderson et al., 2011; Moore et al., 2000; Perani et al., 2001; Wrigley et al., 2009). Despite a lack of compelling evidence in favour of face-to-hand remapping, the assumption of drastic reorganisation in humans remains highly influential in the neuroscientific literature and the clinic (Kaas et al., 2008; Kandel, 2013). Reorganisation of affected cortical areas is thought to drive both recovery of function and the formation of maladaptive neuronal circuitry that relates to neuropathic pain (Jutzeler, Curt, & Kramer, 2015; Moxon, Oliviero, Aguilar, & Foffani, 2014; Zaaimi, Edgley, Soteropoulos, & Baker, 2012; Zörner et al., 2014). Given this proposed role of S1 reorganisation in determining clinical outcomes, it is essential to determine whether the human S1 cortex does, in fact, undergo large-scale reorganisation after an SCI.

Several factors may explain the inconsistency of face-to-hand area remapping results in humans compared to non-human primates. Firstly, studies in non-human primates have primarily reported chin-to-hand area reorganisation in S1 (Jain et al., 1997; Jain, Qi, Collins, & Kaas, 2008; Kambi et al., 2014; Pons et al., 1991), whereas human studies have focused on mouth-to-hand area reorganisation (Corbetta et al., 2002; Henderson et al., 2011; Wrigley et al., 2009). If cortical reorganisation is limited to regions neighbouring the deprived hand area, then differences in facial somatotopy between non-human primates and humans may account for the lack of mouth-to-hand area reorganisation. Indeed, the S1 face representation is inverted in non-human primates, with the chin neighbouring the hand representation (Kaas, 1983). In contrast, human studies show that the forehead is positioned closest to the hand representation (Kikkert, Sonar, Freund, Paik, & Wenderoth, 2023; Penfield & Boldrey, 1937; Root et al., 2022). Secondly, there is no methodological consensus in the literature on how best to quantify reorganisation in humans. Experimental approaches are varied and typically limited to non-invasive techniques such as fMRI, which are not directly comparable to electrophysiological experiments employed in animal models. Notably, while tactile stimulation protocols have been employed in non-human primates, human studies often use movement paradigms due to the difficulty of providing tactile face stimulation in an MRI environment (Corbetta et al., 2002; Curt et al., 2002; Lotze, Laubis-Herrmann, Topka, Erb, & Grodd, 1999; Mikulis et al., 2002; Turner et al., 2001). Moreover, most studies quantify reorganisation using Euclidean distance measures (Henderson et al., 2011; Turner et al., 2001; Turner et al., 2003; Wrigley et al., 2009), which do not account for the folded anatomy of the cortical surface. Alternatively, geodesic distances would provide a more anatomically and functionally accurate measure of the spatial relationship between hand and face cortical representations (Oligschläger et al., 2017). Lastly, researchers studying human reorganisation have classically relied on univariate analysis techniques. However, multivariate methods may be more effective than univariate approaches for studying cortical remapping, as they evaluate the entire multidimensional activity pattern elicited by stimuli, enabling the detection of subtle representational changes that may not emerge as differences in overall activity levels. To date, only Root et al. (2022) have utilised a multivariate representational similarity analysis (RSA) approach to study reorganisation in this context. These methodological inconsistencies highlight the need for more systematic and sensitive approaches to assess reorganisation in human fMRI studies.

In this study, we conducted an in-depth investigation to reassess the assumption of face-to-hand area remapping in the human S1 cortex following SCI using fMRI and extensive uni-and multivariate analysis techniques. In Experiment 1, we replicated previous human fMRI studies using a lip movement paradigm to characterise mouth-to-hand area reorganisation. In Experiment 2, drawing on insights from non-human primate work, we applied vibrotactile stimulation of the forehead, lips, and chin to characterise S1 face-to-hand reorganisation in detail. By stimulating multiple face parts, we ensured that the cortical neighbour of the hand area was not overlooked. Moreover, for a comprehensive analysis, we assessed face-to-hand reorganisation using both univariate and multivariate analysis approaches. We hypothesised that if face representations reorganise into the S1 hand area in human cervical SCI patients, we would find increased face activity in the S1 hand area. Alternatively, if face representations shift toward but do not fully invade the deprived S1 hand area, we would detect a shift in the activity elicited by lip movements and/or face stimulation in SCI patients relative to controls. Furthermore, we expected to find more distinct face information in the S1 hand area, reflected by higher representational dissimilarity scores in SCI patients compared to controls. Given the greater sensitivity of multivariate approaches to detect subtle changes, we further explored correlations with these results to better understand what clinical and behavioural determinants may drive face-to-hand reorganisation in the human somatosensory cortex. By combining several measures of reorganisation, we aimed to provide a systematic and robust framework to assess face-to-hand reorganisation in human cervical SCI patients.

## 2. Materials and methods

### 2.1 Participants

Seventeen chronic (i.e., >6 months post-injury) cervical SCI patients and twenty-one age-sex- and handedness-matched able-bodied control participants (mean age ± s.e.m. = 49.9 ± 3.4; 2 females; 3 left-handers) participated in this study. Patient inclusion criteria were as follows: Aged 18-75 years, no MRI contraindications, at least 6 months post cervical SCI, no neurological impairment or body function impairments not induced by SCI, and able to provide informed consent. One patient was excluded after data collection upon confirming their injury was thoracic, not cervical, resulting in a final sample of sixteen cervical spinal cord injury patients (mean age ± s.e.m. = 52.4 ± 3.5 years; 1 female, 3 left-handers; years since injury ranging from 6 months to 37 years; see Table 1). There was no significant difference in mean age between groups (t_(34.08)_ = 0.53, p = 0.6029; two-tailed).

**Table 1:** Patient demographic and clinical details. Sex: M = male, F = female; Age = age in years; Time since injury = Time since injury in years; AIS grade = American Spinal Injury Association (ASIA) impairment Scale grade defined based on the International Standards for Neurological Classification of Spinal Cord Injury (ISNCSCI): A = complete, B = sensory incomplete, C = motor incomplete, D = motor incomplete; Neurological level of injury = defined based on the ISNCSCI; GRASSP score = Graded Redefined Assessment of Strength, Sensibility and Prehension (able-bodied score: 232 points). Patients are colour-coded according to time since injury, consistent across all figures.

|  | Sex<br>(M/F) | Age<br>(YRS) | Time since injury<br>(YRS) | AIS grade<br>(A/B/C/D) | Neurological<br>level of injury | GRASSP score |
| --- | --- | --- | --- | --- | --- | --- |
| PT01 | M | 42 | 0.5 | D | C3 | 188 |
| PT02 | M | 59 | 1 | D | C1 | 165 |
| PT03 | M | 38 | 1 | D | C4 | 159 |
| PT04 | M | 52 | 1 | D | C7 | 168 |
| PT05 | M | 44 | 4 | D | C4 | 154 |
| PT06 | M | 35 | 7 | A | C4 | 22 |
| PT07 | F | 70 | 7 | D | C5 | 153 |
| PT08 | M | 38 | 7 | A | C4 | 77 |
| PT09 | M | 69 | 9 | D | C2 | 164 |
| PT10 | M | 31 | 11 | C | C5 | 133 |
| PT11 | M | 55 | 13 | D | C4 | 142 |
| PT12 | M | 74 | 19 | D | C7 | 148 |
| PT13 | M | 43 | 21 | A | C6 | 85 |
| PT14 | M | 70 | 29 | A | C4 | 99 |
| PT15 | M | 60 | 36 | C | C5 | 136 |
| PT16 | M | 59 | 37 | A | C7 | 166 |

Control participant inclusion criteria were as follows: Aged 18-75 years, no MRI contraindications, no impairment of body function induced by SCI, no neurological illness, no hand impairments, and able to provide informed consent. The data of one control participant was excluded from the analysis of experiment 1 due to corruption of their fMRI data during data acquisition. Ethical approval was granted by the Kantonale Ethikkommission Zürich (KEK-2018-00937), and written informed consent was obtained prior to the study onset. This study is registered on clinicaltrials.gov under the number NCT03772548. A portion of the control participant dataset used in Experiment 2 has been previously published (see Kikkert et al., 2023 for details).

### 2.2 Clinical characterisation

Behavioural testing was conducted in a separate session. We classified each patient’s injury completeness and impairment level using the International Standards for Neurological Classification of Spinal Cord Injury (ISNCSCI). In addition, we used the Graded Redefined Assessment of Strength, Sensibility and Prehension (GRASSP) assessment to define the sensory and motor integrity of the upper limbs (Kalsi-Ryan et al., 2012). We determined each patient’s most impaired upper limb according to the maximum GRASSP score for each limb. Finally, to assess pain, all SCI patients completed the European Multicenter Study About Spinal Cord Injury (EMSCI) pain questionnaire (version 4.2, emsci.org/). The average pain intensity over the past week was scored using an 11-point numeric rating scale, with 0 indicating no pain and 10 indicating the worst imaginable pain.

### 2.3 Experimental design

#### Experiment 1

In experiment 1, we investigated face-to-hand area remapping in S1 using a lip movement paradigm. We designed a blocked fMRI experiment with three self-paced movement conditions: (1) face movement (continuous jaw clenching and self-paced lip pursing), (2) left-hand open-and-closing, and (3) right-hand open-and-closing. Note that the hand movement conditions were not included in the analysis of the current manuscript (see Howell et al., 2024 for details).

We instructed participants to focus on a green fixation cross on a black screen throughout the scan. Participants viewed the screen via a mirror mounted on the head coil with the instructions ‘left hand’, ‘right hand’, ‘face’, and ‘rest’ displayed in white text behind the green fixation cross. Each movement block lasted 12 seconds, followed by a 6-second rest block. Each condition was repeated 8 times per run in a counterbalanced order. In total, we acquired four runs, each with a different block order, resulting in a total duration of 28 minutes and 48 seconds. Instructions were delivered using Psychtoolbox (v3) implemented in Matlab (v2014b). Head motion was minimised using over-ear MRI-safe headphones or padded cushions.

#### Experiment 2

In experiment 2, we investigated face-to-hand area remapping using a vibrotactile face stimulation paradigm. We used a newly developed MRI-compatible tactile stimulation device that employs soft pneumatic actuator (SPA) technology to provide focal, suprathreshold vibrotactile stimulation to specific parts of the face. We previously described our device setup and stimulation paradigm in Kikkert et al. (2023). However, for accessibility, we reiterate the stimulation setup below.

We used a soft, skin-like interface that uses modulated pneumatic pressure to provide vibrotactile stimulation. The soft pneumatic actuators (SPA) skins are made of MR-safe silicone (Dragon Skin 30®, Smooth On Inc., USA), and the amplitude is controlled by varying the internal pressure as well as its actuation frequencies (Sonar, Gerratt, Lacour, & Paik, 2019; Sonar & Paik, 2016). The SPA system consists of the following components: a stimulus computer, a portable air compressor, a portable pneumatic setup, and pneumatic tubes. We controlled stimulation intensity and frequency using a stimulus computer in the scanner control room. An air compressor, also placed in the scanner control room, provided airflow to a pressure regulator that was controlled by a microcontroller in the portable pneumatic setup. The stimulation intensity could be adjusted using the pressure regulator and an array of high-speed pneumatic valves that were driven by high-power switches and inputs from the microcontroller. A 50 mm diameter hole in the RF shielding wall then carried the 5–6 m long and 4 mm thick tubes (2.8 mm I.D.) to the SPA-skin in the fMRI scanner room. The entire SPA setup is optimised for meeting the fMRI specifications for the selection of individual components, as discussed in detail by Joshi, Sonar, and Paik (2021).

We attached ring-shaped SPA-skins to the forehead (∼1cm above the eyebrow), above the upper lip, and on the chin (Figure 2A). We selected these sites to ensure that an SPA-skin was placed on each of the face areas innervated by the three trigeminal nerve branches. We took special care not to place SPA-skins on the areas that lie at the border of trigeminal nerve innervations. To ensure good grounding of the SPAs on the face, we placed in-house 3D printed plates on top of the SPA-skins and used a custom-made fabric face mask to apply light pressure to the SPA-skins. A further SPA-skin was attached to the ipsilateral thumb using adhesive tape. This condition was part of the broader stimulation protocol and was not included in the current analysis. Instead, we focus on cortical representations of intact body parts, i.e., the face, to avoid confounding input level differences with representational changes. Notably, differences in cortical thumb activity may reflect impaired communication between the periphery and the CNS, rather than differences in the representation itself, which may be maintained even in the absence of peripheral somatosensory input (Kikkert et al., 2021). Moreover, in the SCI group, higher stimulation intensities were applied to the thumb, making direct comparisons with the control group methodologically inappropriate.

Participants viewed a visual display positioned at the head of the scanner bore through a mirror mounted on the head coil. To cue the participant which location would be stimulated, a white text cue (“forehead”, “lips”, “chin”, and “thumb”) appeared on a black screen 0.8s prior to stimulation onset and remained on the screen until stimulation offset. Participants were instructed to attend to the highlighted stimulation location while the cue was displayed. Stimulation was presented for 8s at 8Hz, consisting of 400ms bursts of stimulation ‘on’ periods followed by a 100ms ‘off’ period to minimise peripheral adaptation. To ensure stable attention during the fMRI runs, stimulations were interrupted in a small percentage of the stimulation blocks (10-20% of blocks per run). In these interrupted stimulation trials, stimulation was provided for 4s, after which a 2s silent period was introduced, followed by another 2s of stimulation. We ensured that the interrupted stimulation trials were equally distributed across the stimulation locations within each run. Participants were instructed to count the number of interrupted stimulation trials and verbally report this at the end of each run.

To ensure that the BOLD response induced by stimulation was not confounded by differences in stimulation intensity across the stimulation locations, we aimed to match sensation intensity across stimulation locations prior to the fMRI runs. We performed a stimulation intensity matching task in which participants first determined the optimal stimulation intensity for the forehead. The forehead was selected as the reference location because it is innervated by the fewest mechanoreceptors and was, therefore, assumed to be the least sensitive stimulation location (Corniani & Saal, 2020). Participants were instructed to identify an optimal stimulation intensity that was as strong as possible while remaining focal (i.e., localised without spread to adjacent skin), comfortable, and stable throughout the 8s stimulation period (i.e., with minimal peripheral adaptation). Participants used a button box to indicate whether the intensity should be increased, decreased, or left unchanged. Based on these responses, the air pressure provided to the SPAs was adjusted in steps of 5 kPa to increase or decrease stimulation intensity or remain stable. If a participant indicated twice in succession that the stimulation intensity should not change, the pressure for the forehead was set for all subsequent fMRI tasks. The current SPA-skins design allows for a pressure input range of 20-80 kPa while producing 0.1-1 N force feedback, suitable for the human tactile perception range (Choi & Kuchenbecker, 2013; Sonar, Huang, & Paik, 2021). Once the optimal stimulation intensity was set for the reference location, participants were asked to match the stimulation intensity for the other stimulation locations to that of the forehead. To facilitate this matching, participants were initially given 8s of stimulation on the forehead, immediately followed by stimulation of one of the other locations. Participants were instructed to adjust the intensity of the second location to match the intensity of the reference location as closely as possible. As before, if the participant indicated twice that the stimulation intensity should remain unchanged, the pressure for that stimulation location was set at this level for all fMRI tasks.

To uncover face somatotopy, we used a blocked design consisting of five conditions: stimulation of the forehead, lips, chin, and thumb, as well as a rest condition (no stimulation). Within each participant, we consistently tested either the right side of the face and right thumb or the left side of the face and left thumb. The side tested was always ipsilateral to the most impaired side of the body for each SCI patient. We then matched the tested sides in controls to those in patients. The visual cue during the face stimulation blocks was as described above and the presentation of the word “Rest” indicated the rest condition. Each condition had a block duration of 8.8s and was repeated eight times per run in a counterbalanced order. Each run comprised a different block order and had a duration of 6min and 7.4s. We acquired four blocked design runs, with a total duration of 24min and 29.6s. Instructions and stimulations were delivered using Psychtoolbox (v3) implemented in Matlab (v2014b). Matlab communicated with the Arduino board implemented in the SPA controller set-up via a serial port over a proprietary protocol. Head motion was minimised using over-ear MRI-safe headphones or padded cushions.

### 2.4 MRI acquisition

We acquired MRI data with a 3 Tesla Philips Ingenia system (Best, The Netherlands) and a 15-channel HeadSpine coil. However, in the instance that the 15-channel coil was unsuitable due to head size limitations, a 32-channel HeadSpine coil was used instead (i.e., PT02, PT03, PT08, PT08, PT09, PT11). We acquired an anatomical T1-weighted structural scan with the following acquisition parameters: TE: 4.4ms; TR: 9.3ms; flip angle: 8°; resolution: 0.75mm^3^. For task-fMRI, we used an echo-planar-imaging (EPI) sequence with partial brain coverage.

For experiment 1, we centred 32 sagittal aligned parallel to the floor of the fourth ventricle, with coverage over S1, the thalamus, and the brainstem. The following sequence parameters were used: single-shot gradient echo, echo time (TE): 26ms; repetition time (TR): 2500ms; flip angle: 82°; spatial resolution: 1.45 x 1.8 x 1.45mm^3^; field of view (FOV): 210 x 187.6 x 70.2mm; SENSE factor: 2.1; with 179 volumes per run). To facilitate post hoc physiological noise correction of the fMRI data, we recorded chest movement data using a respiratory bellow throughout the scans (sampling rate: 496Hz). A pulse oximeter was attached to the second digit of each participant’s foot to record oxygen saturation and cardiac pulse. All physiological measurement devices were connected to a data acquisition device (Invivo; Phillips Medical Systems, Orlando, Florida) coupled with a desktop computer running the corresponding recording software.

For experiment 2, we centred 36 sagittal over the postcentral gyrus, with coverage over the thalamus and brainstem. The following sequence parameters were used: single-shot gradient echo, echo time (TE): 30ms; repetition time (TR): 2200ms; flip angle: 82°; spatial resolution: 2.3mm^3^; field of view (FOV): 210 x 210 x 82.8mm; SENSE factor: 2.1; with 167 volumes per run. We additionally acquired sagittal T2w structural images of patients’ cervical spinal cord at the lesion level to quantify spared tissue bridges. These scans were acquired in a separate testing session using a 3 Tesla MAGNETOM Prisma Siemens Healthcare system (Erlangen, Germany) with a 64-channel head/neck coil. The sagittal T2w sequence was acquired using the following parameters: TR: 3500ms; TE=84ms; flip angle: 160; echo train length: 14; spatial resolution: 0.3x0.3x2.5mm; FOV 220mm x 220mm x 55mm.

### 2.5 MRI preprocessing and analysis

fMRI data preprocessing was applied to each individual fMRI run using FSL’s Expert Analysis Tool FEAT (version 6.0; https://fsl.fmrib.ox.ac.uk/fsl/fslwiki/FEAT). To ensure consistent alignment of the hemisphere of interest in Experiment 1, we flipped the raw data along the x-axis for participants whose most impaired body side was the right side (or matched for controls). Similarly, in Experiment 2, we applied the same procedure for participants for whom the right side of the face was tested. This ensured that the hemisphere contralateral to the most impaired side of the body (in Experiment 1) or the tested face side (in Experiment 2) was consistently aligned.

#### Experiment 1

As experiment 1 was part of a larger study that focuses on brainstem imaging (Howell et al., 2024), we applied an optimised brainstem fMRI preprocessing pipeline based on previous research (Beissner, 2015; Beissner, Deichmann, & Baudrexel, 2011; Faull, Jenkinson, Clare, & Pattinson, 2015; Matt et al., 2019; Meissner et al., 2024). This pipeline included brain extraction using FSL’s automated brain extraction tool (BET (Smith, 2002)), motion correction using FMRIB’s Linear Image Registration Tool (MCFLIRT (Jenkinson, Bannister, Brady, & Smith, 2002)), a 90s high-pass temporal filter implemented via FSL’s Expert Analysis Tool (FEAT (Woolrich, Ripley, Brady, & Smith, 2001)), and spatial smoothing using a 3mm full-width-at-half-maximum (FWHM) isotropic Gaussian kernel. Further denoising was completed using FSL’s Physiological Noise Modelling (PNM (Brooks et al., 2008)). Cardiac and respiratory phases were assigned slicewise to each volume of the EPI image time series. The physiological noise regression model comprised a total of 34 regressors: 4th-order harmonics for both the cardiac and respiratory cycles, 2nd-order harmonics for their interactions, and individual regressors for heart rate and respiration volume. We visually inspected the waveforms to ensure accurate identification of the peaks of the cardiac and respiratory cycles, making adjustments as needed. The resulting voxelwise confound regressors were incorporated into the first-level general linear model (GLM).

Independent component analysis (ICA) was conducted using the Multivariate Exploratory Linear Optimized Decomposition interface (MELODIC (Smith et al., 2004)). The resulting components were classified as either “Noise” or “Signal” based on visual inspection of their spatial maps, time series, and power spectra information, following the guidelines published by Griffanti et al. (2017). Two independent researchers labelled the components to assess interrater reliability. Cohen’s kappa indicated strong agreement between raters, with an estimated score of 0.85 (95% CI: 0.82 to 0.88). The ICA-derived noise regressors were subsequently added to the GLM design matrix.

First-level parameter estimates were computed using a voxel-based general linear model (GLM) based on the gamma hemodynamic response function and its temporal derivatives. Time series statistical analysis was conducted using FILM (FMRIB’s Improved Linear Model) with local autocorrelation correction. Confound regressors were created for the hand movement tasks. Face movement was contrasted with rest. A fixed effects higher-level analysis was run for each participant to average across runs.

#### Experiment 2

Standard preprocessing steps were included for the experiment, including brain extraction using an automated brain extraction tool BET (Smith, 2002), motion correction using MCFLIRT (Jenkinson et al., 2002), spatial smoothing using a 3mm full-width-at-half-maximum (FWHM) Gaussian kernel, and high-pass temporal filtering using a cut-off of 90s. First-level parameter estimates were computed using a voxel-based general linear model (GLM) based on the gamma hemodynamic response function and its temporal derivatives. Time series statistical analysis was conducted using FILM (FMRIB’s Improved Linear Model) with local autocorrelation correction. To reduce noise artefacts, scan-wise cerebrospinal fluid (CSF) and white matter time series were added to the model as nuisance regressors. Contrasts were defined for each face stimulation condition versus rest. A further regressor of no interest was defined for the 0.8s in which a visual cue was provided without tactile stimulation. A fixed effects higher-level analysis was run for each participant to average across runs.

#### 2.5.1 MRI co-registration

We registered each participant’s fMRI data to the T1-weighted image, using 6 degrees of freedom and the mutual information cost function, and then optimised using boundary-based registration (BBR (Greve & Fischl, 2009)).

#### 2.5.2 Region of interest definition

Face-to-hand area remapping was studied using anatomical regions of interest (ROIs) of S1, obtained from an atlas (Glasser et al., 2016), and the S1 hand area (obtained via https://github.com/DiedrichsenLab/fs_LR_32). We further segmented this S1 ROI into 50 bands (Figure 1A), as in Schone et al. (2023), to study the distribution of activity across this region.

**Figure 1:**
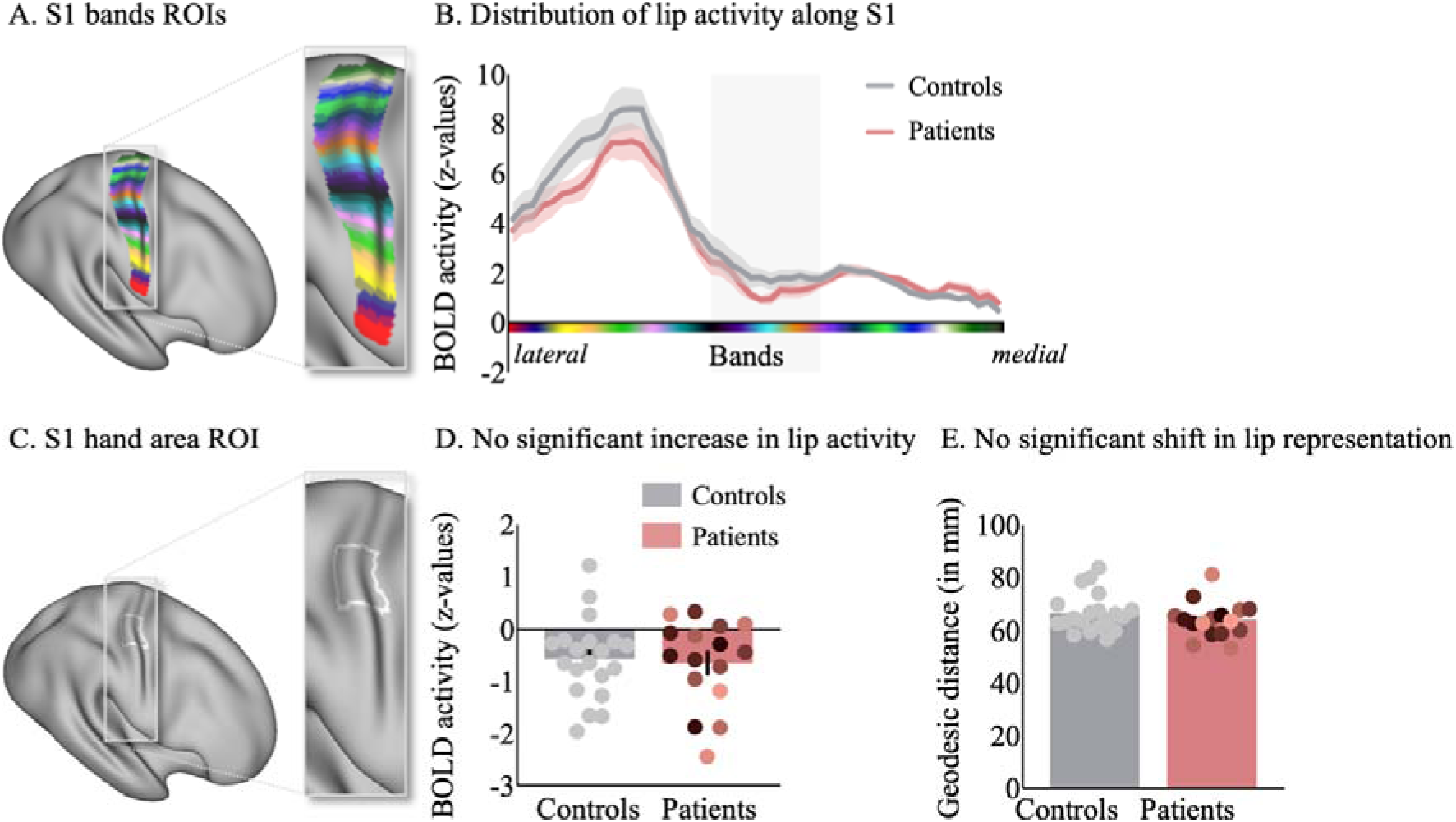
No significant evidence in favour of face-to-hand area remapping using a lip movement paradigm. A) S1 region of interest (ROI) divided into 50 distinct bands. B) Lip movement successfully activated S1 in both controls (grey) and patients (red). Shaded areas represent the standard error of the mean. Peak activity during lip movement was highest in the lateral S1 bands. The grey shaded bands represent the bands that overlap with the S1 hand area ROI in C). D) There was no significant difference in lip movement activity in the S1 hand area between patients (red) and controls (grey). Dots represent individual participants with patients colour-coded according to time since injury, as annotated in Table 1. Error bars represent the standard error of the mean. E) We calculated the geodesic distance in mm for each participant between an anchor in the S1 foot area and peak lip movement activity. Geodesic distances between peak lip activity and the S1 anchor did not differ significantly between groups.

We used an anatomical mask of the S1 hand area that has previously been used to investigate somatotopic hand representations and plasticity (Beukema, Diedrichsen, & Verstynen, 2019; Kikkert et al., 2016; Kikkert, Pfyffer, Verling, Freund, & Wenderoth, 2021; Kolasinski et al., 2016; Wesselink et al., 2022). The S1 hand area ROI was further restricted by only selecting the nodes 1 cm below and above the anatomical hand knob within the S1 area ROI (i.e., the midpoint of the S1 hand area mask; Figure 1C). We selected this conservative boundary of the hand ROI to ensure that our ROI did not expand into the S1 face area. The S1 hand area ROIs were first projected onto the individual participant’s inflated cortical surface and then onto their 3D T1-weighted images. The cortical surface projections were constructed from each participant’s T1-weighted images using FreeSurfer v6.0 (https://surfer.nmr.mgh.harvard.edu/) (Dale, Fischl, & Sereno, 1999; Fischl, Liu, & Dale, 2001) and Connectome Workbench v1.3.2 (https://www.humanconnectome.org/software) (Marcus et al., 2011).

#### 2.5.3 Functional MRI analysis

##### S1 bands analysis

To investigate the distribution of face activity across S1, the statistical maps from the second-level analysis of experiments 1 and 2 were projected onto each participant’s cortical surface using cortical-ribbon mapping and then onto the fs-LR surface template brain. We then extracted the peak activity (z-value) in each of the 50 S1 ROI bands.

##### Level of activity analysis

To explore univariate activity, we extracted the average z-value under the S1 hand area ROI contralateral to the most impaired body side during lip movement versus rest (experiment 1) or contralateral to the stimulated face side during forehead, lip, or chin stimulation versus rest (experiment 2).

##### Cortical distance analysis

To assess for a shift in face activity in SCI patients relative to controls, we extracted the peak vertex (i.e., maximally activated vertex on the FS-LR inflated surface) for each participant during lip movement (experiment 1) and for each of the stimulation conditions (experiment 2) within the S1 area ROI. We then calculated the geodesic distance from an anchor placed in the S1 foot area (vertex 5550 on the fs-LR surface template brain) to the peak vertex for forehead, lip and chin stimulation.

##### Representational similarity analysis

To gain insights beyond the broad topographic distribution of the face representation, we employed multivariate representational similarity analysis (RSA). This method allows us to capture more subtle changes in the relationships between face-evoked activity patterns within the S1 hand area. Unlike traditional univariate analyses, RSA considers the full multidimensional activity pattern elicited by tactile stimulation of a body part, making it more sensitive to detecting underlying sensorimotor representations. We, therefore, calculated the dissimilarity between the activity patterns for each pair of face stimulation conditions within the S1 hand area.

We estimated the representational dissimilarity between face locations using the cross-validated squared Mahalanobis distance (Nili et al., 2014; Wesselink & Maimon-Mor, 2017), adhering to previously established procedures (Kikkert et al., 2021; Odermatt et al., 2024; Root et al., 2022; Wesselink et al., 2019). We first extracted the voxel-wise parameter estimates (betas) and the model fit residuals under the ROI, then prewhitened the betas using the model fit residuals. Next, we calculated the cross-validated squared Mahalanobis distances between each pair of conditions, using the four runs as independent cross-validation folds. The resulting distances were averaged across the folds. The dissimilarity values for all pairs of conditions were initially assembled in a representational dissimilarity matrix (RDM), with a width and height corresponding to the number of conditions. To quantify the strength of the representation or “representational separability”, we averaged the unique off-diagonal values of the RDM. Here, if conditions cannot be statistically differentiated (i.e. when a parameter is not represented in the ROI), the expected value of the distance estimate is 0. In contrast, if activity patterns can be distinguished, this value will be larger than 0.

#### 2.5.4 Structural MRI analysis

##### Midsagittal tissue bridge analysis

We used sagittal T2w structural images of the cervical spinal cord at the lesion level to quantify spared tissue bridges (Huber, Lachappelle, Sutter, Curt, & Freund, 2017; Pfyffer, Huber, Sutter, Curt, & Freund, 2019; Vallotton et al., 2019). The segmentation was performed using Jim 7.0 software (Xinapse Systems, Aldwincle, UK), which has demonstrated high intra- and interobserver reliability (Huber et al., 2017; Pfyffer et al., 2019). The experimenter conducting the manual segmentation was blinded to patient identity. Images from two patients were excluded due to metal artefacts or insufficient data quality, which would prevent reliable lesion quantification. Tissue bridges were defined as the relatively hypointense intramedullary region between the hyperintense CSF on one side and the cystic cavity on the other. We measured the smallest width of ventral and dorsal tissue bridges on the midsagittal slice and summed them to obtain the total width of the remaining tissue bridges.

### 2.6 Statistical analysis

We carried out statistical analysis using JASP (v0.17.2.1). We used standard approaches for statistical analysis, as mentioned in the Results section. Outliers were defined as values that were more than 3 standard deviations greater or lower than the group mean and were replaced with group-specific averages.

If normality was violated (assessed using the Shapiro-Wilk test), appropriate non-parametric statistical testing was used. All group testing was two-tailed. We corrected for multiple corrections per reorganisation measure in our group comparisons using the Bonferroni correction. Uncorrected p-values are reported with the adjusted alpha-levels. Given the exploratory nature of our correlational analysis, no correction for multiple comparisons was carried out here.

Bayesian analysis was conducted for the main comparisons to investigate support for the null hypothesis with the Cauchy prior width set at 0.707 (i.e., JASP’s default). If assumptions of normality were violated, we ran group comparisons using Mann Whitney Bayesian testing with 1000 permutations. Following the conventional cut-offs, a Bayes Factor (BF) smaller than 1/3 is considered substantial evidence in favour of the null hypothesis. A BF greater than 3 is considered substantial evidence and a BF greater than 10 is considered strong evidence in favour of the alternative hypothesis. A BF between 1/3 and 3 is considered weak or anecdotal evidence (Dienes, 2014; Kass & Raftery, 1995).

For the S1 bands analysis, comparisons were made between SCI patient and control data for each band using non-parametric permutation t-tests in the SPM1D Matlab toolbox for one-dimensional data (SPM1D M.0.4.12; https://spm1d.org/). SPM1D utilises random field theory (RFT) to make statistical inferences on one-dimensional data by treating discrete data points, such as individual bands, as part of a continuous field. This approach accounts for spatial correlations in the data and estimates the probability of observing a supra-threshold test statistic under the null hypothesis.

## 3. Results

### Experiment 1: Investigating face-to-hand area remapping using a movement paradigm

In Experiment 1, we aimed to replicate the procedures from previous studies using a lip movement paradigm to investigate face-to-hand area remapping in S1. First, we visualised the distribution of lip movement activity along S1 by extracting the peak lip movement activity in each of our S1 bands (Figure 1A-B). Consistent with the somatotopic organisation of the somatosensory homunculus, we saw the highest lip movement activity in the lateral bands of S1 and lower activity in the medial bands of S1 (Catani, 2017; Penfield & Boldrey, 1937). Notably, we did not find significant differences in the distribution of face activity across these bands between SCI patients and control participants (non-parametric permutation t-tests; largest Z= -2.05, *P*LJ= 0.37; smallest Z = 0.03, *P* > 0.99).

We further hypothesised that face-to-hand area remapping in cervical SCI patients would result in increased activity in the S1 hand area (Figure 1C) during lip movements. However, group differences were non-significant (t_(34)=_0.28, p = 0.78; Figure 1D), and the Bayes factor (BF10 = 0.33) indicated anecdotal to moderate evidence in favour of the null hypothesis. Crucially, if the lip representation shifted towards but did not invade the S1 hand area, then it is possible that no changes in lip movement activity levels would be observed in the S1 hand area, but rather, there would be a smaller geodesic distance between peak lip activity and an anchor placed in the S1 foot area. However, no significant difference in lip-to-anchor distance between SCI patients and controls was found (t_(34)_ = 1.10, p = 0.28; BF = 0.52; Figure 1B), with the Bayes factor suggesting anecdotal evidence in favour of the null hypothesis. Similar results were found in S1 area 3b (see Supplementary Figure 3).

### Experiment 2: Investigating face-to-hand area remapping using a tactile stimulation paradigm

In experiment 2, we used a tactile face stimulation paradigm. Since the lip representation does not neighbour the hand representation in humans, we expanded our investigation to include the forehead, lips, and chin, ensuring that each skin area innervated by the distinct trigeminal nerve branches was represented in our investigation (Figure 2A). We found that our custom-built device, based on soft pneumatic actuator (SPA) technology, effectively activated S1 (Figure 2B-D). A qualitative inspection of the distribution of peak activity along S1 showed that while lip and chin stimulation predominantly activated S1 lateral of the S1 hand area, forehead stimulation appeared to be represented both lateral and medial to the S1 hand area. We did not find significant differences in the distribution of face activity across these bands between SCI patients and control participants for the forehead (largest Z= - 2.15, *P* = 0.37; smallest Z = -0.01, *P* > 0.99), lip (largest Z= 2.13, *P* = 0.38; smallest Z = - 0.03, *P* > 0.99), or chin stimulation (largest Z= -1.82, *P* = 0.53; smallest Z = 0.12, *P* > 0.99). For individual control and patient activation maps of forehead, lip and chin stimulation, see Supplementary Figures 1 and 2.

**Figure 2:**
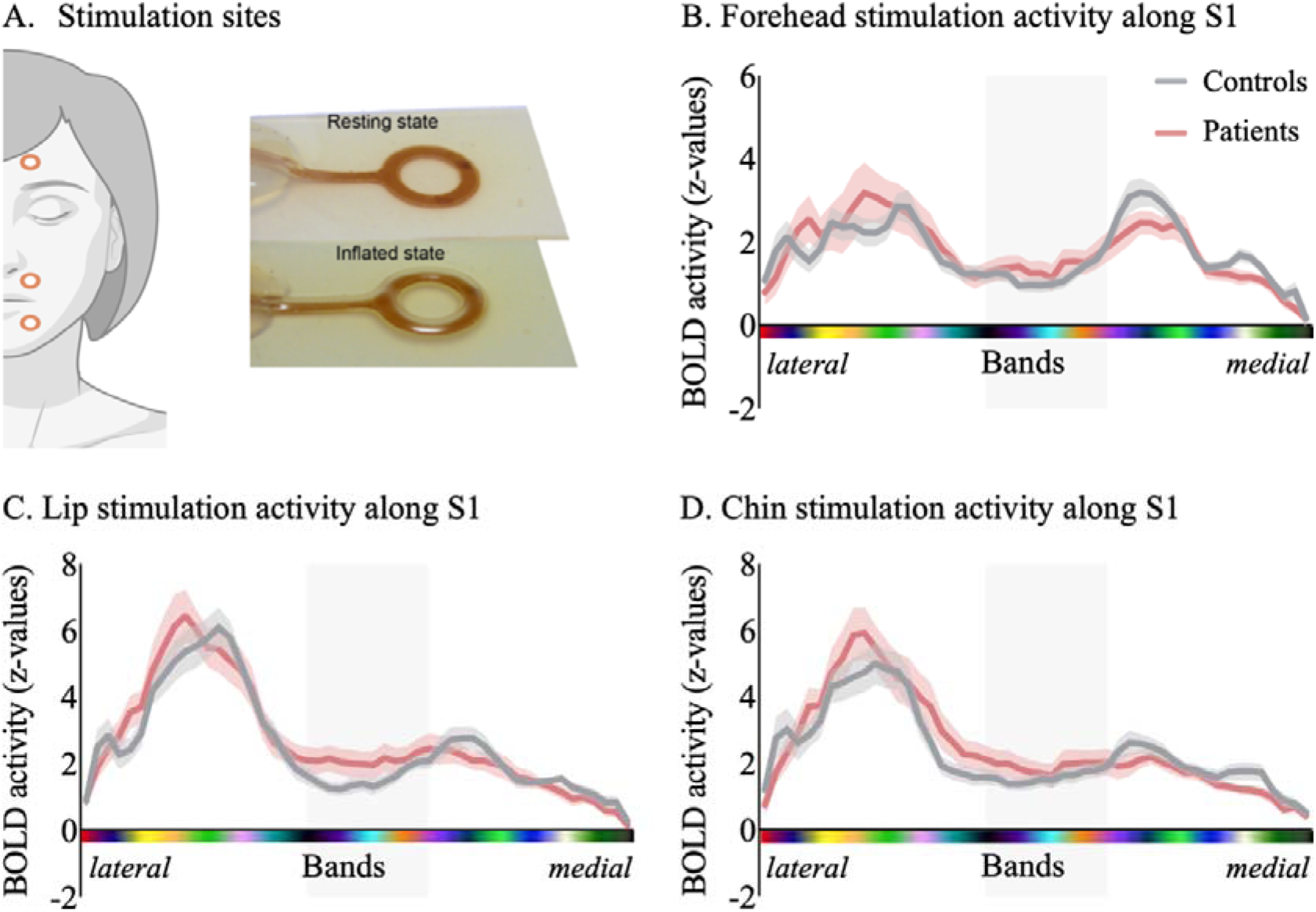
Tactile face stimulation successfully activated S1. A) We applied vibrotactile stimulation to the forehead, lips, and chin using a custom-built device based on soft pneumatic actuator (SPA) technology to study face-to-hand area reorganisation in S1. Peak activity along the S1 band ROIs during stimulation of the forehead (B), lips (C) and chin (D). While peak activity was most predominant in the lateral bands of S1 for lip and chin stimulation, forehead activity appears to be represented both lateral and medial to the S1 hand area. The grey shaded areas represent the bands that overlap with the S1 hand area. Annotations are as in Figure 1.

As in Experiment 1, we hypothesised that if large-scale face-to-hand area remapping had occurred in cervical SCI patients, we would observe increased activity in the S1 hand area (Figure 3A) and/or a shift in peak activity (Figure 3B) in patients compared to controls during forehead, lip, or chin stimulation. However, we did not find a significant difference in SCI patients compared to controls during stimulation of the forehead (t_(35)_ = 0.70, p = 0.49; BF_10_ = 0.39; adjusted α = 0.017), lips (t_(35)_ = 0.41, p = 0.69; BF_10_ = 0.34), or chin (t_(35)_ = 0.23, p = 0.82; BF_10_ = 0.33), with Bayes factors suggesting anecdotal evidence for the null hypothesis. We similarly did not observe a significant shift in peak activity during forehead (t_(35)_ = -0.86, p = 0.40, BF_10_ = 0.44; adjusted α = 0.017), lips (t_(35)_ = -0.24, p = 0.81, BF_10_ = 0.34), or chin stimulation (W = 169, p = 0.37, BF_10_ = 0.43) between groups. Notably, forehead peak activation showed a bimodal distribution across participants. Therefore, to ensure that a potential shift in peak forehead activity was not obscured by the broader distribution across the sensorimotor cortex, we repeated the analysis after restricting the S1 ROI to the S1 hand area and the inferior S1 region, increasing sensitivity to lateral-to-medial shifts. The restricted analysis did not reveal statistically significant group effects (i.e., S1 face area; forehead: (W = 212, p = 0.18; BF_10_ = 0.74; adjusted α = 0.017; lip: W = 143.5, p = 0.46, BF_10_ = 0.39; chin: t_(35)_ = -0.55, p = 0.59, BF_10_ = 0.36).

**Figure 3:**
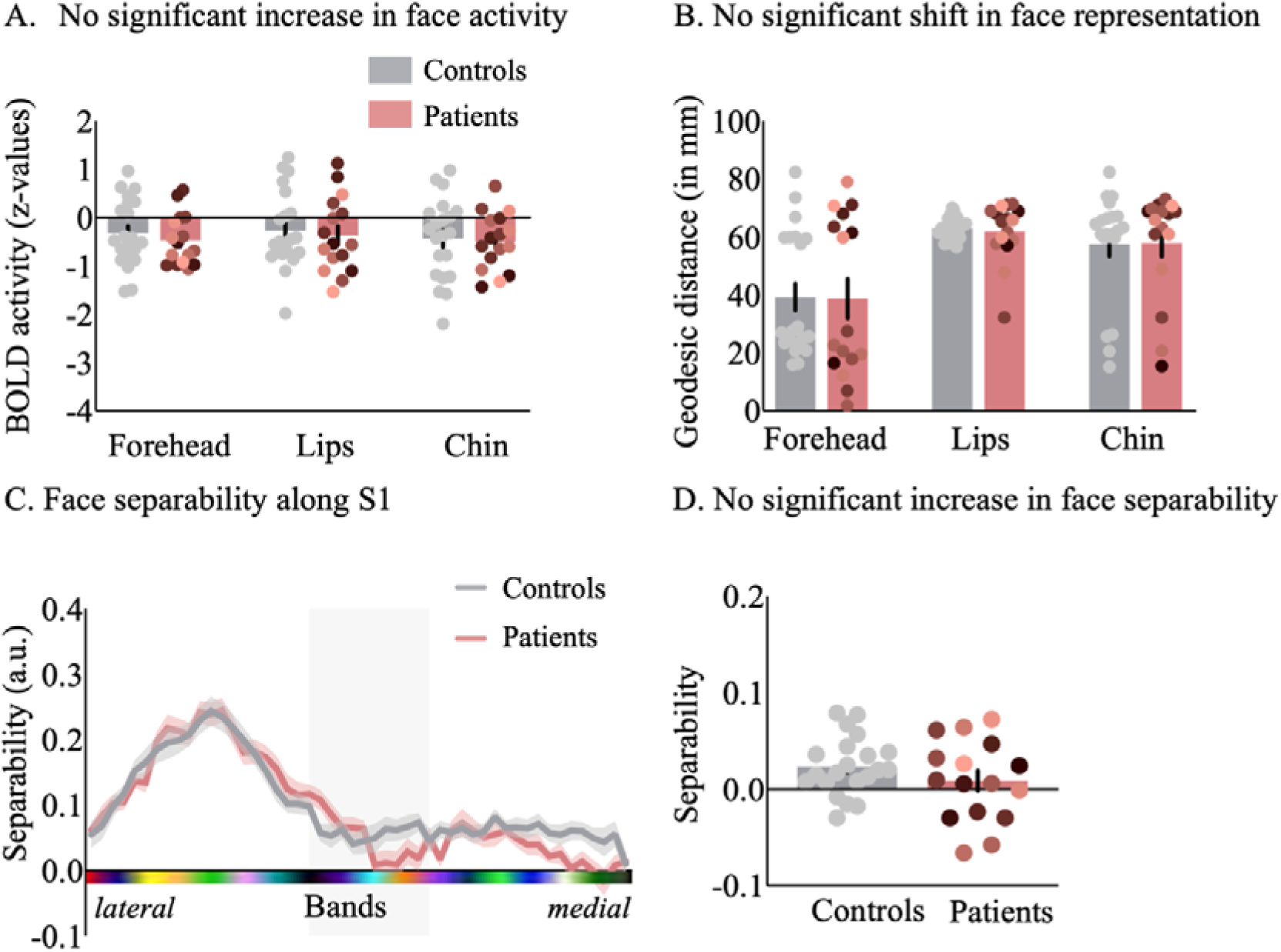
A tactile stimulation paradigm did not show significant evidence in favour of face-to-hand area reorganisation in S1. A) Average activity in the S1 hand area during forehead, lip, and chin stimulation did not differ significantly between patients and controls. B) We calculated the geodesic distance in mm for each participant between an anchor in the S1 foot area and peak activity for each of our stimulation conditions. Geodesic distances between peak forehead, lip and chin activity and the S1 anchor did not differ significantly between groups. C) Mean face-part separability scores along the S1 band ROIs. The grey shaded area*s* represent the bands that overlap with the S1 hand area. D) A comparison of face-part separability within the S1 hand area did not reveal significant differences between patients and controls. Annotations are as in Figure 2.

Next, we used multivariate representational similarity analysis (RSA) to gain insights into the underlying activity pattern elicited by tactile stimulation of the face. We hypothesised that if face-to-hand reorganisation had occurred, there would be more distinct information in the S1 hand area, allowing for better differentiation between the face conditions. In other words, the average representational separability (or ‘representation strength’) between the activity patterns elicited by stimulating different face parts would be higher. First, we visualised the separability, measured as the mean inter-face parts distance, across each of our S1 bands for both the SCI patients and controls (Figure 3C). We saw greater representational separability in the bands lateral to the S1 hand area, with no significant difference in separability scores between the SCI patient and control groups across the S1 band ROIs (largest Z= 2.83, *P* = 0.10; smallest Z = 0.02, *P* > 0.99). A comparison of face-part separability within the S1 hand area revealed no significant difference in representational separability in cervical SCI patients compared to controls (Figure 3D; W = 133, p = 0.45; BF_10_ = 0.41), with the Bayes factor showing anecdotal results in favour of the null hypothesis.

### Clinical correlates of face-to-hand remapping

Next, we aimed to understand which clinical, behavioural, and structural spinal cord determinants may drive face-to-hand area remapping in S1 of cervical SCI patients. Given that we assume the representational similarity analysis is the most sensitive in detecting reorganisation, we focused our correlational analysis on our outcome measure of face-to-face part representational separability in the S1 hand area (Figure 4). We did not find any significant correlations with the number of years since injury (r_s_ = -0.23, p = 0.38), retained sensorimotor function of the upper limbs (GRASSP; r*_s_* = 0.34, p = 0.20), pain (r*_s_* = -0.01, p = 0.98), or spared midsagittal spinal tissue bridges at the lesion level (r*_s_*= 0.08, p = 0.79).

**Figure 4:**
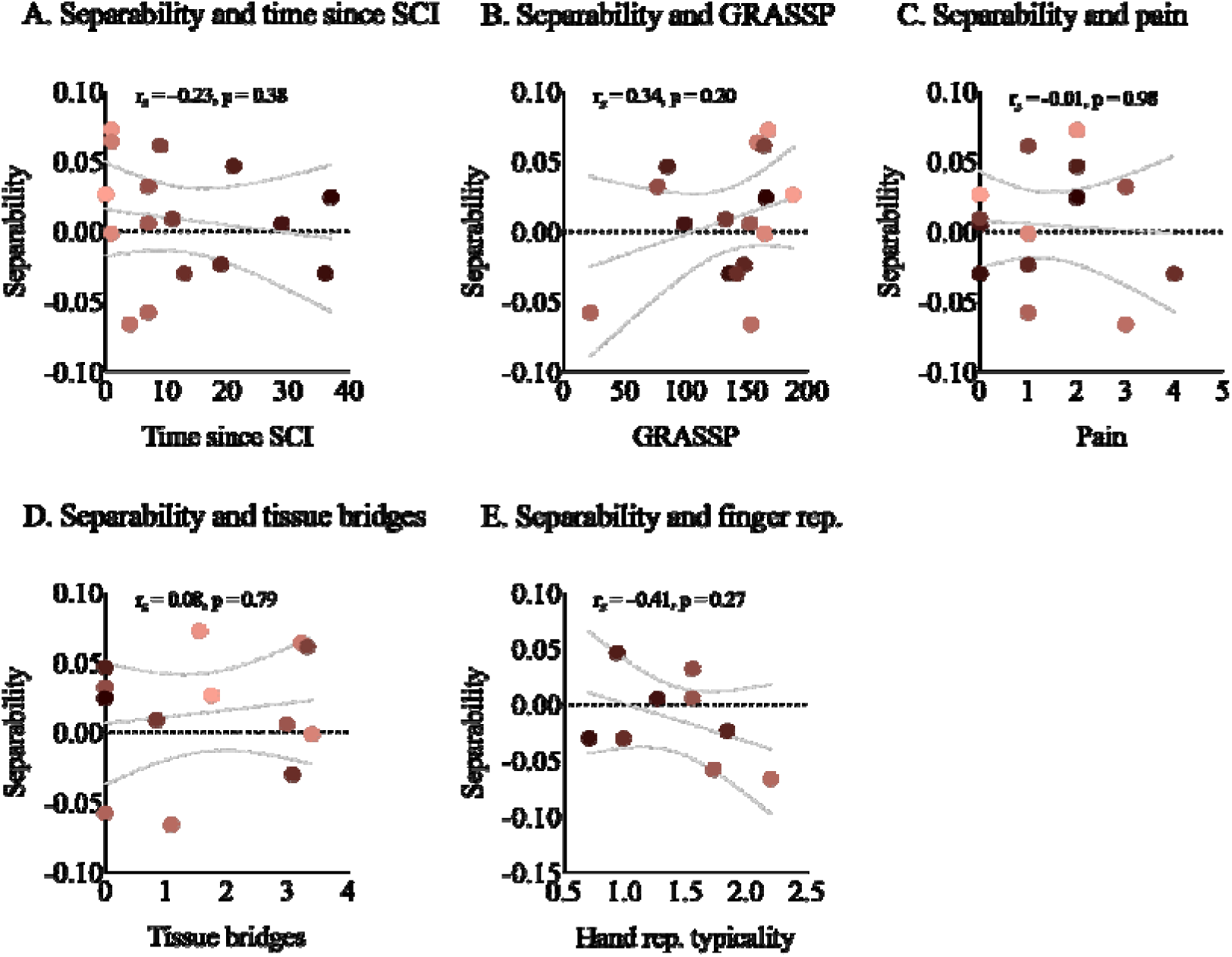
No significant correlations between S1 face-to-hand area remapping and clinical, behavioural, and structural spinal cord measures. We examined clinical, behavioural, and spinal cord structural correlates for face-to-hand area remapping, quantified using mean face-part representational separability in the S1 hand area. There was no significant correlation between representational separability and time since SCI (A), sensorimotor function of the upper limbs as measured using the GRASSP (B), pain (C), and spared midsagittal spinal tissue bridges at the lesion level (D). Finally, we found that preservation of the S1 hand representation does not significantly correlate with S1 face-to-hand area remapping (E). Rep. = representation. Annotations are as in Figure 2.

Preservation of the S1 hand representation may interfere with S1 face-to-hand area remapping. We, therefore, explored whether SCI patients with more salient hand representation in the S1 hand area exhibit less face-to-hand area remapping. A subset of patients from this study (n = 9) also participated in our previous study, where we examined preserved finger somatotopy in cervical SCI patients (Kikkert et al., 2021). We extracted the somatotopic hand representation typicality (or normality) for each of these patients from our previous study. However, we did not find a significant correlation (-0.41, p = 0.27), suggesting that the preservation of the S1 hand representation does not influence potential face-to-hand area remapping.

## 4. Discussion

In this study, we aimed to resolve inconsistencies in the human reorganisation literature and assess face-to-hand area remapping in cervical SCI patients in unprecedented detail. To do so, we utilised both movement (experiment 1) and vibrotactile stimulation paradigms (experiment 2), allowing us to investigate the full architecture of S1 face reorganisation. We found no significant evidence in favour of changes in face representations following SCI compared to control participants across three distinct reorganisation measures. Furthermore, we found no significant correlations between face-to-hand area reorganisation and the clinical, behavioural, and structural outcome measures. These results suggest that large-scale face-to-hand area remapping may not be apparent after cervical SCI in humans.

Most prior human research into cortical face reorganisation following SCI has focused exclusively on the lower face, primarily the mouth (Corbetta et al., 2002; Curt et al., 2002; Henderson et al., 2011; Lotze et al., 1999; Mikulis et al., 2002; Turner et al., 2001; Wrigley et al., 2009). Instead, our study considered multiple parts of the face, systematically assessing each of the distinct facial areas innervated by trigeminal nerve branches. Our findings suggests that face representations may be stable in the chronic phase after SCI, with no significant evidence showing remapping of either the upper or lower face areas into the deprived hand region. This adds to the growing body of literature suggesting stable sensorimotor representations in humans following sensory deprivation and extends it to individuals with cervical SCI (Andersen & Aflalo, 2022; Bruurmijn, Pereboom, Vansteensel, Raemaekers, & Ramsey, 2017; Flesher et al., 2016; Kikkert et al., 2016; Kikkert et al., 2021; Makin & Krakauer, 2023; Root et al., 2022; Schone et al., 2023; Wesselink et al., 2022; Wesselink et al., 2019). These preserved representations can be utilised to restore sensorimotor functions through brain-computer interfaces. For instance, through intracortical microstimulation in the S1 hand area, researchers have been able to elicit fine-grained tactile sensations within the hand of cervical SCI patients several years after their injury (Armenta Salas et al., 2018; Flesher et al., 2016; Flesher et al., 2021; Hiremath et al., 2017).

Yet, how can such contrasting accounts of stability and reorganisation coexist in the literature? Through a homeostatic plasticity model, Muret & Makin (2021) propose that body-part representations are more broadly distributed across the homunculus than previously believed. Upon sensory deprivation, these underlying body-part representations are unmasked, which may be misinterpreted as reorganisation. Our finding of two distinct peaks for forehead activation, alongside similar evidence from others (Gordon et al., 2023; Muret, Root, Kieliba, Clode, & Makin, 2022), supports the notion of distributed body-part representations across the sensorimotor cortices. Critically, a limitation of previous reorganisation research, in both animals and humans, is that cortical maps are often defined using a winner-take-all approach (Makin & Krakauer, 2023; Wesselink et al., 2022). Through this approach, any underlying representational overlap between body parts is disregarded, with the boundaries of cortical areas defined according to the dominant body-part response. Therefore, in the absence of hand activity following an SCI, the native S1 hand area may be reassigned to the next dominant body part included in the analysis, i.e. the face, and subsequently misinterpreted as evidence for cortical reorganisation. In the present study, we addressed this methodological confound by avoiding winner-take-all approaches and employing RSA to account for the potential overlap of underlying representations.

Through RSA, we were able to investigate not only whether new face information emerges in the deprived somatosensory cortex, but also whether there is a difference in how this face information is structured. We found no significant group differences in face-part separability in the cortical hand area. Moreover, face-part separability did not show significant associations with clinical or behavioural measures, suggesting that these factors may not strongly influence the organisation of face representations in this region. However, the absence of significant associations should be interpreted with caution, particularly given the reduced sample sizes for the correlations involving spared midsagittal tissue bridges (n = 13) and preserved hand representation typicality scores (n = 9). Across multiple analysis approaches, we found no statistically significant evidence of face-to-hand reorganisation. Yet, Bayesian analyses provided only anecdotal to moderate support for the null hypothesis, which does not warrant definitive conclusions. Future studies would benefit from larger sample sizes to rule out subtle reorganisation effects.

While we used vibrotactile stimulation to target multiple face regions, key methodological limitations remain that may contribute to the discrepancy in results between animal models of SCI and human SCI patients. Fundamentally, animal studies typically model SCI via uni- or bilateral transection of the ascending dorsal column (Jain et al., 2008; Kaas et al., 2008; Massey et al., 2006; Onifer et al., 2005; Reed et al., 2016). In contrast, human SCI typically arises from a traumatic incident, leading to a pattern of disruption that is more anatomically and clinically heterogeneous and often results in both sensory and motor impairments with a range of completeness. This clinical heterogeneity may favour alternative forms of reorganisation, such as the expansion of impaired forearm representations in patients with incomplete SCI, rather than face-to-hand reorganisation. In the present cervical SCI cohort, the face represents the only sensory input that is consistently unimpaired across participants, whereas preserved upper limb function varies substantially (as indexed by GRASSP scores), which restricts a systematic investigation of alternative reorganisation patterns.

Furthermore, we used 3T fMRI, which indirectly measures thousands of neurons within a single voxel, whereas invasive electrophysiological recordings utilised in animal models are capable of detecting neural activity at a single-neuron resolution. While higher-field imaging may provide greater spatial resolution to detect reorganisation, it is typically contraindicated in tetraplegic patients with cervical implants (Hoff et al., 2019). Other modalities, such as resting-state connectivity or EEG, may also capture network-level or temporal changes that we could not resolve with task-based fMRI. Despite this limited resolution, previous fMRI studies in non-human primate models of SCI have been able to demonstrate face reorganisation (Chen, Qi, & Kaas, 2012; Dutta, Kambi, Raghunathan, Khushu, & Jain, 2014). Moreover, fine-grained intracortical stimulation of the deprived S1 hand area in human cervical SCI patients has been shown to elicit sensations only within the paralysed hand and not in adjacent body parts, such as the face. Therefore, it is unlikely that these methodological discrepancies solely account for the absence of face-to-hand remapping in our human SCI cohort.

It is also possible that large-scale face reorganisation, if present in humans, occurs predominantly during the acute phase following injury. Although a substantial body of non-human primate research has demonstrated persistent neural reorganisation in the chronic phase following SCI (in both complete and incomplete injuries; (Dutta et al., 2014; Florence & Kaas, 1995; Jain et al., 1997; Jain, Qi, Raman, Lyon, & Kaas, 2025; Jain et al., 2008; Kambi et al., 2014; Kambi, Tandon, Mohammed, Lazar, & Jain, 2011; Liao, Reed, Kaas, & Qi, 2016; Pons et al., 1991), we did not find evidence supporting reorganisation in the chronic phase in the present study. Accordingly, we cannot exclude the possibility that face-to-hand area reorganisation occurred earlier after injury and stabilised prior to entry into the chronic phase.

Importantly, while our findings do not demonstrate evidence in favour of face-to-hand remapping in human SCI patients, this does not imply that reorganisation may never occur in humans. Reorganisation has been documented in other contexts, such as congenital limb loss (Hahamy et al., 2017; Kamping, Lütkenhöner, & Knecht, 2004; Root et al., 2021; Root et al., 2022; Stoeckel, Seitz, & Buetefisch, 2009). Yet, emerging evidence suggests that this reorganisation is restricted to the prenatal or early developmental stages (Root et al., 2021; Root et al., 2022; Schone et al., 2023; Stoeckel et al., 2009; Tucciarelli et al., 2024). A recent longitudinal study by Schone et al. (2023) demonstrated that the adult human brain is remarkably stable. Specifically, the researchers tested individuals before and after arm amputation and showed that somatosensory representations of the face and missing arm remained unaltered. Thus, while non-human primate research suggests that the somatosensory system of adult mammals remains highly plastic (Jain, Florence, Qi, & Kaas, 2000; Jain et al., 2008; Merzenich et al., 1983; Pons et al., 1991), it is possible that the temporal limits of cortical reorganisation in humans differ.

Together, our findings call for the assumption of large-scale face-to-hand remapping in human SCI patients to be revisited. Specifically, our results emphasise the need to critically reassess the translation of findings from animal models to humans. Further research is necessary to uncover the specific conditions under which large-scale cortical remapping may occur in humans and better understand its potential role in recovery following SCI.

## Data availability

Aggregated data will be made available online upon publication.

## Acknowledgements

We thank the participants for their participation in the study, Sijamini Baskaralingam and Ingrid Odermatt for their help with data collection, Roger Luchinger and the Institute for Biomedical Imaging for scanning support.

