## Supplementary Material for "Revisiting face-to-hand area remapping in the human primary somatosensory cortex after a cervical spinal cord injury"

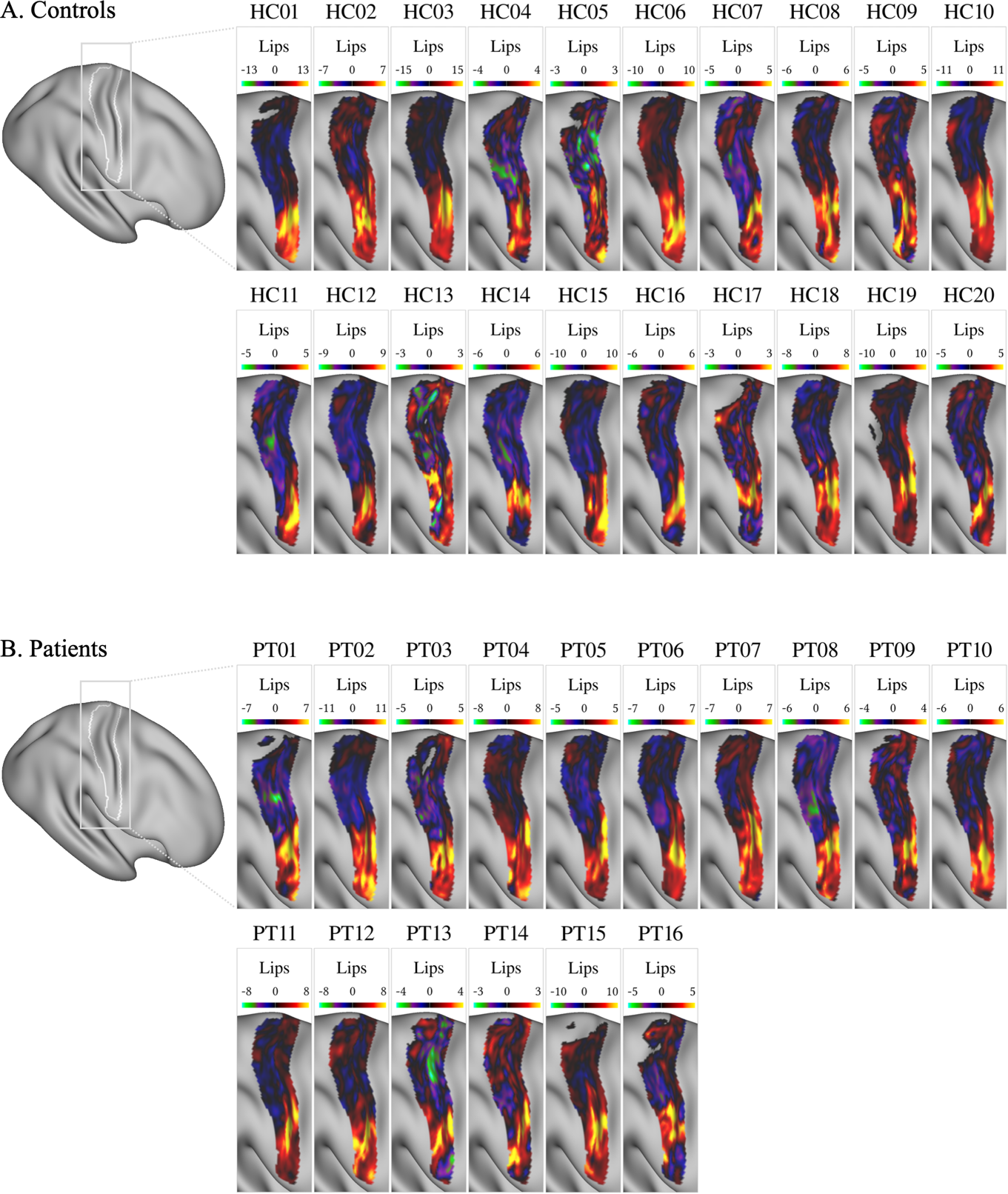

**Supplementary Figure 1: Activation maps in the primary somatosensory cortex during lip movement.** Individual activation maps for control participants (A) and spinal cord injury patients (B) showing the contrast lips > rest. Maps are displayed on the fs_LR cortical surface and restricted to the primary somatosensory cortex. Colours indicate uncorrected z-statistics, as shown in the corresponding colour bar. Patients are ordered by increasing time since injury.

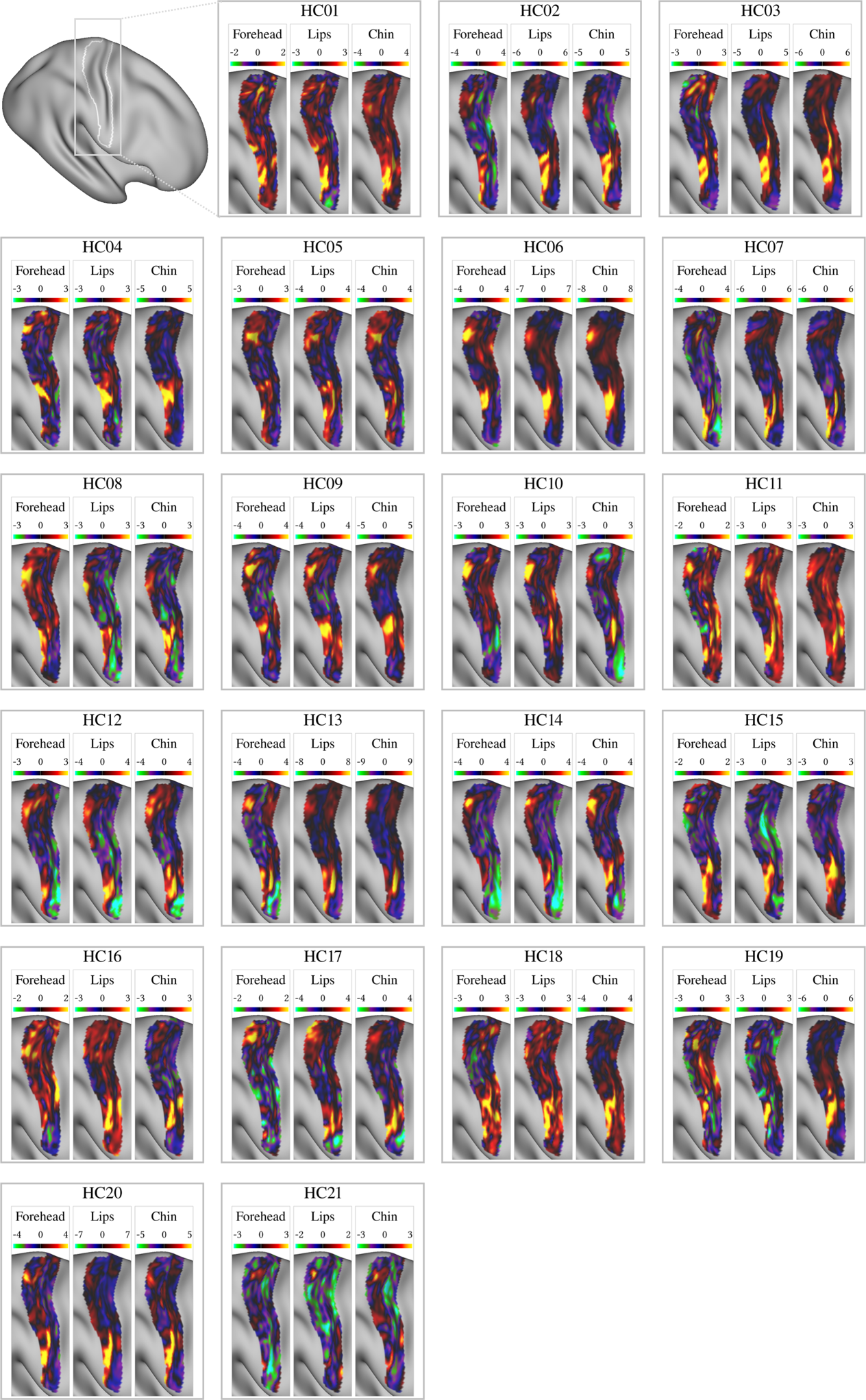

**Supplementary Figure 2: Activation maps in the primary somatosensory cortex of control participants during forehead, lip and chin stimulation.** Individual activation maps for control participants showing the contrasts forehead > rest, lips > rest and chin > rest. Maps are displayed on the fs_LR cortical surface and restricted to the primary somatosensory cortex. Colours indicate uncorrected z-statistics, as shown in the corresponding colour bar.

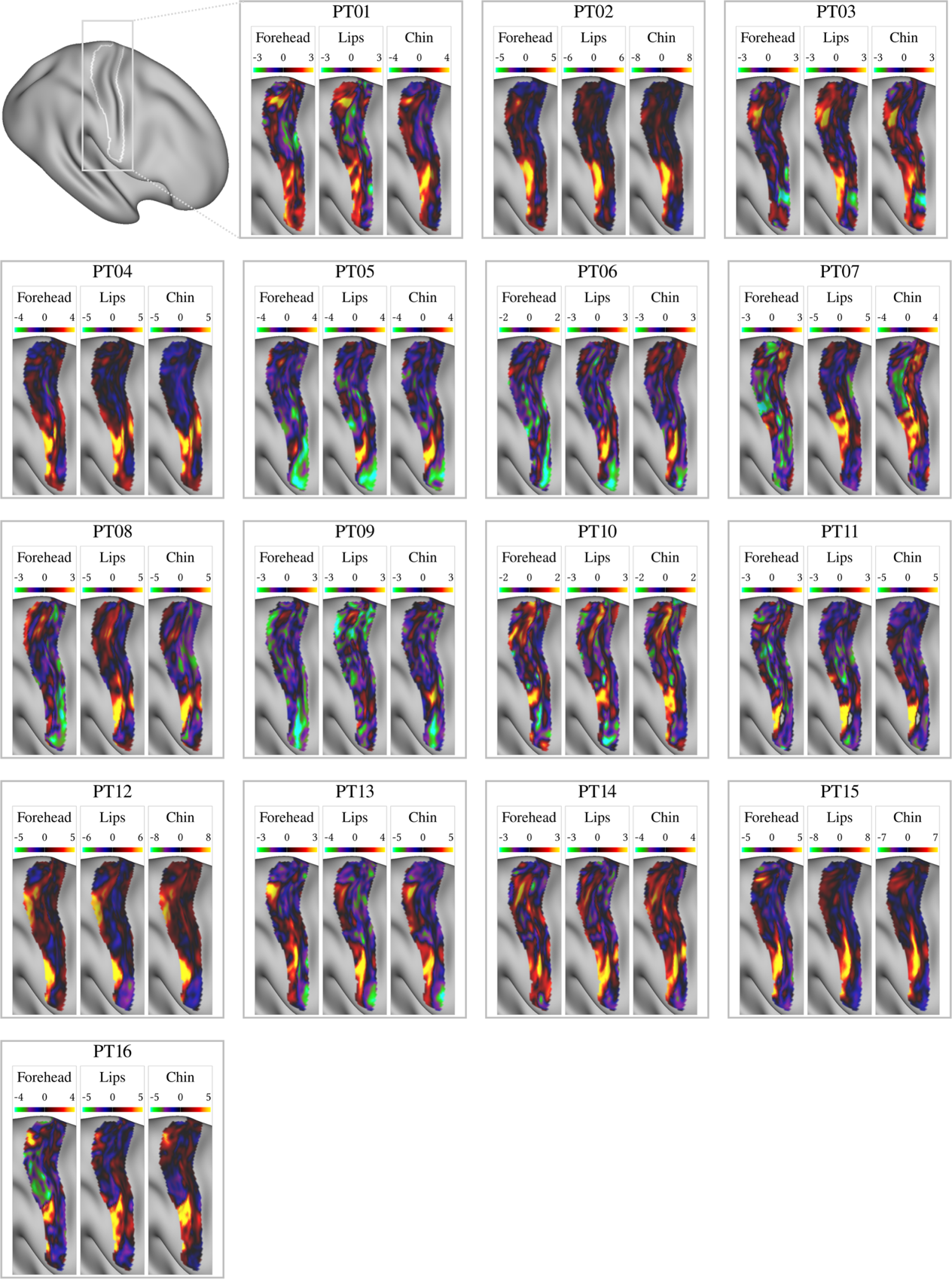

**Supplementary Figure 3: Activation maps in the primary somatosensory cortex of SCI patients during forehead, lip and chin stimulation.** Individual activation maps for spinal cord injury patients showing the contrasts forehead > rest, lips > rest and chin > rest. Maps are displayed on the fs_LR cortical surface and restricted to the primary somatosensory cortex. Colours indicate uncorrected z-statistics, as shown in the corresponding colour bar. Patients are ordered by increasing time since injury.
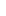
